# Genomic imprinting in an early-diverging angiosperm reveals ancient mechanisms for seed initiation in flowering plants

**DOI:** 10.1101/2024.09.18.613646

**Authors:** Ana M. Florez-Rueda, Mathias Scharmann, Leonardo P. de Souza, Alisdair R. Fernie, Julien B. Bachelier, Duarte D. Figueiredo

## Abstract

The evolution of the seed habit marks a pivotal innovation of the spermatophytes. Angiosperms further refined this trait by coupling the development of seed accessory structures to fertilization, optimizing resource allocation. Here, we demonstrate that post-fertilization auxin production is an evolutionarily conserved mechanism for seed initiation in angiosperms. We also provide evidence that this pathway likely emerged from a switch from maternal to paternal control after the divergence of angiosperms from their gymnosperm ancestors. Our study thus brings new insights into the evolutionary origins of the endosperm, which was a determining feature for the rapid rise to dominance of flowering plants.

## Main text

The plesiomorphic seed habit of spermatophytes is the most successful method of sexual reproduction in vascular plants. But there are key differences between the seeds of its two clades, gymnosperms and angiosperms. In gymnosperm seeds, the embryo is the only biparental structure and is nourished by a haploid maternal tissue derived from the proliferation of the megagametophyte^1,2^. In angiosperms, however, the development of the seed nourishing structure, the endosperm, is tightly linked to fertilization. This is because in angiosperms both the embryo and the endosperm are biparental, and result from the fertilization of two maternal gametes, egg cell and the central cell, by two paternal sperm cells. These two fertilization products interact with each other and the formation of the endosperm is indispensable for embryo viability^3^. Moreover, these structures are surrounded by a seed coat of maternal sporophytic tissues, which provides various functions^4^, and whose development is dependent on signals originating in the biparental endosperm^5–7^. This coupling to fertilization has obvious advantages, as it prevents the allocation of nutrients to “empty seeds”, which do not carry an embryo, and is therefore a major innovation of angiosperms.

Using the model dicot species *Arabidopsis thaliana* (Arabidopsis), we previously demonstrated that formation of the biparental endosperm is linked to the paternal expression of auxin biosynthesis genes^8^. This phenomenon, known as genomic imprinting, involves the parent-of-origin specific allele expression of certain genes^9^. The main auxin biosynthesis pathway is a two-step process, the first being catalyzed by TRYPTOPHAN AMINOTRANSFERASES (TAA/TARs) and the second by YUCCA (YUC) flavin monooxygenases^10^. Before fertilization, the expression of genes encoding these enzymes is repressed in the maternal gametophyte by H3K27me3 repressive marks^8,11^, which prevents auxin production in the absence of fertilization. However, the paternal alleles of *TAA/TAR* and *YUC* genes are not labeled with these repressive epigenetic marks, and are thus expressed in the fertilized central cell^8^. This means that upon gamete fusion, auxin biosynthesis initiates in a paternal-dependent fashion, triggering the precocious development of the endosperm and of the surrounding sporophytic tissues^5,8^. Consistently, exogenous applications or ectopic production of auxin in unfertilized ovules leads to the formation of asexual endosperms and seed coats, also known as autonomous seeds^5,8^.

Interestingly, genes involved in auxin biosynthesis are imprinted in the endosperm of other eudicots and of monocots^12–14^, and auxin biosynthesis seems to be a hallmark of seed nourishing structures^15,16^. But the degree to which these mechanisms are conserved remains to be assessed. Here, we asked if the imprinting of auxin biosynthesis is evolutionarily conserved in angiosperms and found that, indeed, auxin drives seed development in early-diverging angiosperms which diverged circa 120-150 million years ago from the mesangiosperms (which include the monocot and eudicot clades)^17^. We propose that these seed initiation pathways were present in the gymnosperms and became sexualized and paternally expressed in the angiosperm’s most recent common ancestor.

To test the hypothesis that auxin biosynthesis is an evolutionarily conserved mechanism for seed initiation, we generated the endosperm imprintome of the early-diverging water lily species *Nymphaea caerulea* (order Nymphaeales, **Fig. 1a, Extended Data Fig. S1**). If the mechanisms of seed initiation are the same in early-diverging angiosperms and in mesangiosperms, then these mechanisms most likely coupled seed development to fertilization at the origin of angiosperms. We used low-coverage genome sequencing to screen individuals of *N. caerulea* at the Botanical Gardens of Berlin (BO) and Potsdam, and identified two diploid self-incompatible individuals with a significant level of DNA polymorphisms (henceforth referred to as plant 1 and plant 2; **Extended Data Fig. S1 and Supplementary Table 1**). We used these individuals for reciprocal crosses and collection of endosperm material using laser capture microdissection. Sequencing of endosperm-purified RNAseq libraries allowed us to estimate allele-specific expression for 5441 genes in the *N. caerulea* genome, for both directions of the reciprocal crosses (**Fig. 1b, Supplementary Tables 1 and 2**). The mean inferred maternal proportions were 0.5192 for plant 1 and 0.4814 for plant 2 (**Supplementary Table 1**). Very close to 0.5, as expected for a diploid endosperm^18^. We thus identified 144 candidate Maternally Expressed Genes (MEGs), as those that exceed our imposed threshold of 0.75 maternal proportion in both directions of the cross (**Fig. 1 b-c**, **Supplementary Tables 1 and 2**). Symmetrically, we inferred 170 Paternally Expressed Genes (PEGs), which show <0.25 maternal proportion in both cross directions (**Fig. 1 b-c, Supplementary Tables 1 and 2**). Of the 314 imprinted genes identified in the *N. caerulea* endosperm, we found 57 to be imprinted in at least one other species whose endosperm had been examined for imprinting (**Fig. 1d**). There was an average of seven commonly imprinted genes between species, which is expected as imprinting has been shown to evolve rapidly^14^. Interestingly, we found comparatively more PEGs compared to a recent study on *Nymphaea* imprintomes^19^. There are, however, two main differences between the two studies which may account for this discrepancy: 1) our analysis was done in an intra-specific cross, rather than interspecific; and 2) our endosperm samples were collected at an earlier time point. Because the endosperm in *Nymphaea* seeds is relatively small and devoid of nutrients, and the seed nourishing function is carried out by a maternal sporophytic tissue (i.e., perisperm)^20^, it is reasonable to expect that in waterlilies, paternal effects are increasingly limited as the seed develops. In such a situation, PEGs would only be detectable at earlier stages of endosperm development in *Nymphaea*, as we show here.

**Figure 1.**
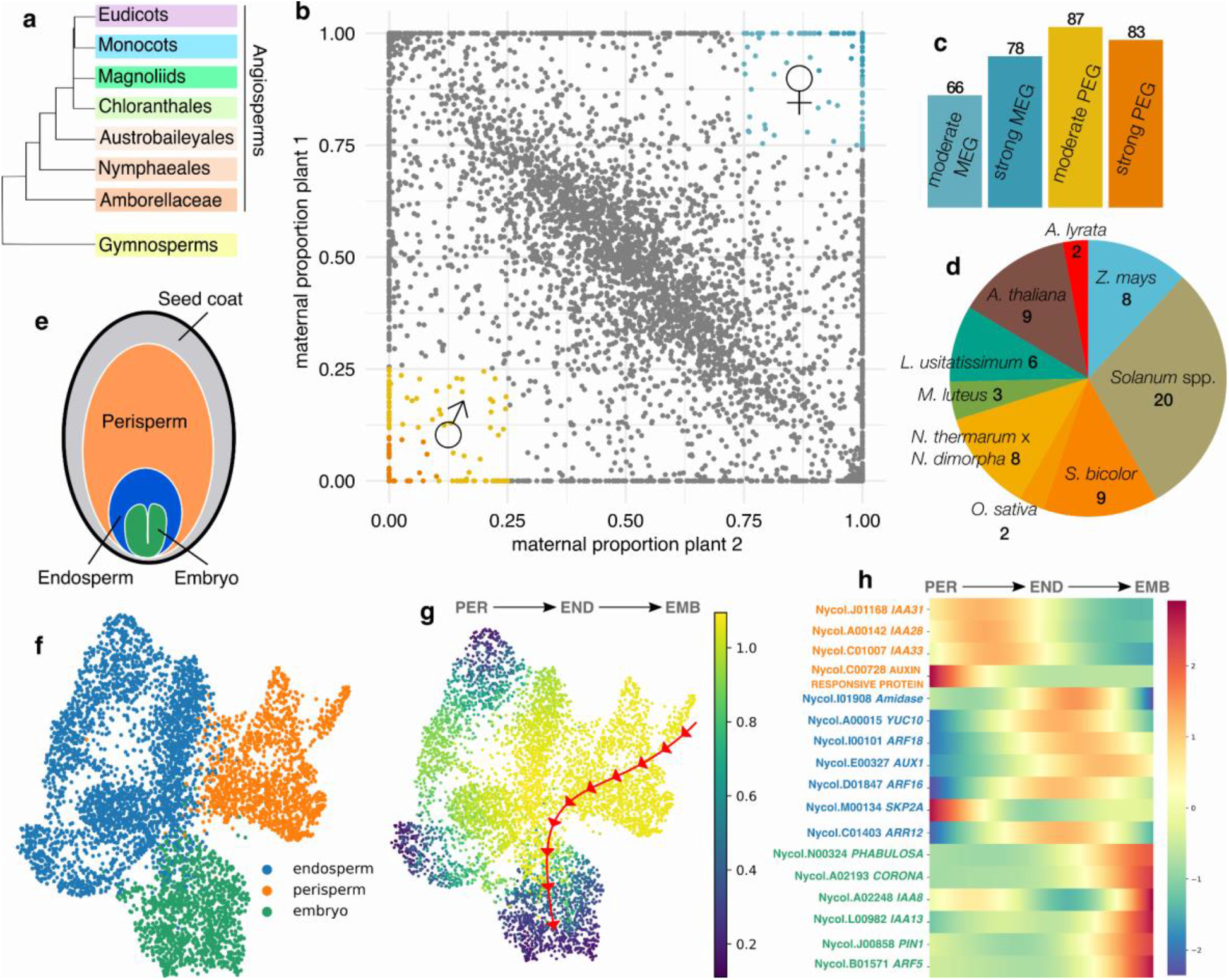
Imprinting and gene expression dynamics in *N. caerulea* endosperm and seed compartments. **a**. Simplified phylogeny of Spermatophytes. **b**. Maternal proportions of 5,441 genes in reciprocal crosses of *N. caerulea* endosperm. Paternally expressed imprinted genes (PEGs, < 25% maternal proportion) are shown in blue, and maternally expressed genes (MEGs, > 75% maternal proportion) in orange. Darker shades indicate strong PEGs/MEGs (PEGs < 10% or MEGs > 90% maternal proportion). **c**. Counts of PEGs and MEGs in *N. caerulea* endosperm, categorized as shown in (b). **d**. Number of commonly imprinted genes shared between *N. caerulea* and other angiosperms. **e**. Diagram of seed compartments in *Nymphaea*. **f**. UMAP visualization of all sequenced nuclei, color-coded by clusters corresponding to different seed tissues as in (e). **g**. Expression trajectory that spans from the perisperm through the endosperm to the embryo. **h**. Heatmaps showing dynamic expression of auxin-related genes along the trajectory outlined in (g).

We then asked which gene ontology (GO) terms were enriched in the set of imprinted genes in *N. caerulea* (**Extended Data Fig. S2**). We found enriched GO terms pointing to pathways with known roles in seed development, including those involved in epigenetic processes (**Supplementary Table 2**). Predicted protein interaction networks can be found in **Extended Data Fig. S2**. Notably, genes involved in hormonal metabolism were also found in enriched functional categories, and this included genes involved in auxin biosynthesis, such as *YUC10* (**Supplementary Table 2**). In fact, the PEG *YUC10* is one of the only three genes that we found imprinted in endosperms of both *N. caerulea* and species of monocots and eudicots (**Supplementary Table 2**), fitting with our hypothesis that paternal expression of auxin biosynthesis genes is conserved. To validate that auxin-related processes are enriched during seed development in *Nymphaea*, we also performed single cell transcriptomics of seed tissues of *N. careulea* and were able to annotate three distinct clusters as embryo, endosperm and perisperm/seed coat (**Fig. 1e and f**) based on reference laser-captured microdissected transcriptomes of the seed tissues^15^**(Extended Data Figs. S3 and S4, Supplementary Table 3**). The single-nucleus analysis demonstrated that the auxin signaling apparatus is subject to a dynamic compartmentalisation across the seed. Specifically, we uncovered a sequential turnover of auxin-related gene expression, progressing from the perisperm to the endosperm, and finally to the embryo **(Fig. 1g, Figure S5)**. The perisperm exhibited high expression levels of the auxin-responsive genes *IAA21, IAA28* and *IAA33*. The endosperm, where the auxin biosynthesis PEG *YUC10* exhibited peak expression, also demonstrated elevated expression of key auxin signaling components, including *ARFs 16* and *18, AUX1*, and *ARR12*. Within the embryo, crucial auxin signaling genes such as *IAA8* and *13, ARF5, PIN1, PHABULOSA*, and *CORONA* were prominently expressed **(Fig. 1f, Figure S5 and S6)**. These findings highlight the critical role of auxin signaling in seed development in *Nymphaea*, providing a detailed map of the spatial compartmentalisation of this signaling pathway at the single-nucleus level.

To validate these findings, we did quantification of active auxin, indole-3-acetic acid (IAA), in somatic and seed tissues of several species of *Nymphaea* (**Extended Data Fig. S7**). While early developing seeds showed IAA levels similar to those of somatic tissues, later stages of seed development coincided with a hyperaccumulation of auxin (**Extended Data Fig. S7**). We also confirmed these observations in giant water lilies (*Victoria cruziana* and *V. amazonica*; **Extended Data Fig. S7**), suggesting that this feature is conserved in the Nymphaeaceae. In fact, auxin accumulation in developing seeds was shown in maize over 80 years ago^21,22^, and this phenomenon has been confirmed in multiple species, both eudicots and monocots^8,23–26^, solidifying that post-fertilization auxin accumulation is a feature of angiosperm seeds.

To further challenge our hypothesis that auxin is an evolutionarily conserved driver of seed development, we tested if exogenous application of auxin is sufficient to bypass pollination and lead to fertilization-independent seed formation in early-diverging lineages. Indeed, unfertilized *N. careulea* ovules treated with auxin increase in size, even quicker than their fertilized counterparts (**Fig. 2a-b**), and show signs of tissue differentiation, namely of the sporophytic tissues. This includes the formation of a perisperm and the differentiation of the endothelial seed coat layer (**Fig. 2a**). However, unlike fertilized seeds, these autonomous seeds are short-lived and many collapse 8 days after the treatments (8 DAT). A defined endosperm structure is also not often seen, and by 8 DAT only remnants of it are observed (**Fig. 2a**). We thus took seed size as a marker for auxin-induced seed formation. Consistent with the role of auxin in driving seed development in *N. careulea*, auxin applications show a marked dose-response in driving autonomous seed formation, and applications of inhibitors of auxin biosynthesis^27,28^ blocks the growth of its fertilized seeds in a dose-dependent manner (**Fig. 2b**). To rule out that this role of auxin is specific to *N. careulea*, we repeated the auxin application to four other species of *Nymphaea*, with identical results (**Extended Data Fig. S8**). The same was true for *Victoria sp*. (**Extended Data Fig. S8**). We then expanded our work to *Schisandra chinensis*, which is a species of liana of the Austrobaileyales clade (**Fig. 1a**). Again, we observed that auxin induces autonomous seed formation in *S. chinensis*, which includes expansion of the nucellus and growth of the surrounding sporophytic structures (**Fig. 2c and Extended Data Fig. S8**). This means that the effect of auxin is not specific to the Nymphaeales, but is also true for other early-diverging angiosperm lineages.

**Figure 2.**
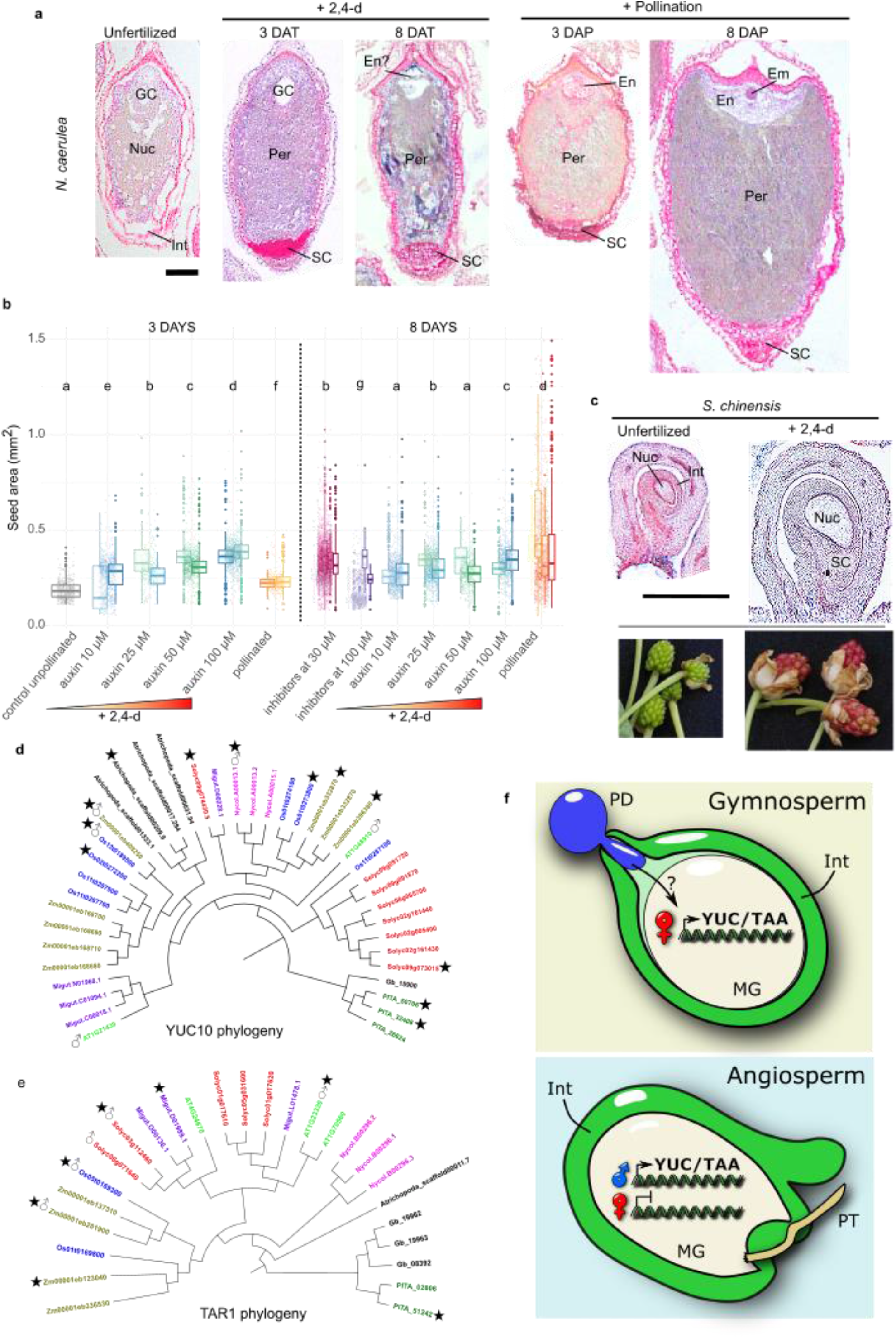
Auxin is an evolutionarily conserved driver of seed development. **a**, Left, unfertilized *N*.*caerulea* ovule. Middle panels, auxin-induced autonomous seeds at 3 and 8 DAT. Right panels, fertilized seeds at 3 and 8 DAP. GC, gametophytic cavity, Int, integuments, Nuc, nucellus, SC, seed coat, Per, perisperm, En, endosperm, Em, embryo. Scale bars, 100 μm. **b**, Size of unfertilized ovules, autonomous and fertilized seeds of *N. careulea*. Left panel, seed sizes at 3 DAT after application of different concentrations of 2,4-D. Right panel, seed sizes at 8 DAT, after application of different concentrations of 2,4-D or of the auxin inhibitors yucasin and L-kynurenine. Each bar represents one fruit and each dot one seed. Letters indicate Wilcoxon rank sum test, P < 0.01. **c**, Top panels, *S. chinensis* ovule (left) and auxin-induced autonomous seed (right) at 10 DAT. Scale bar, 1 mm. Bottom panels, outer ovule morphology in mock and auxin-treated samples. **d-e**, Cladograms of (**d**) Orthogroups OG0001099 (*YUC* orthologs) and (**e**) OG0001584 (*TAR* orthologs) denoting genes enriched in the nourishing tissues of the seed. Orthogroup inference and DGE analyses as inferred previously^15^ and in this study. Imprinted statuses as previously determined^12,13,30–34,37^ and in this study. ★DEGs ♂ PEGs. Dark green, *P. pinaster*, fucsia, *N. caerulea*, bright green: *A. thaliana*, red, tomato, purple *M. lutteus*, blue, *O. sativa*, brown, *Z. mays*, black, other species. **f**, Proposed model of seed initiation by auxin. In gymnosperms, the deposition of pollen in the pollination drop triggers auxin-related processes in the ovule. This coincides with the maternal expression of auxin biosynthesis genes in the megagametophyte. In angiosperms, seed development became coupled to fertilization, and thus auxin biosynthesis in the fertilized central cell switched to paternal control. PD, pollination drop, Int, integuments, MG, megagametophyte, PT, pollen tube.

Our data thus points to post-fertilization auxin biosynthesis being a conserved driver of seed development in the angiosperms. This is supported by the observation that auxin biosynthesis genes are endosperm PEGs in *Nymphaea* (this study) and in species from which the Nymphaeales diverged over 100 Mya ago^8,12,29–34^. This is particularly striking as there is limited conservation in imprinting even among closely related lineages^14,35^, and auxin biosynthesis genes are thus the exception to that trend. We thus postulated that those genes should have a common evolutionary origin and should be in the same clade. Based on our previous comparative transcriptomic dataset of seed nourishing tissues^15^, we identified orthogroups OG0001099 and OG0001584, which contain the land plant *YUC* and *TAR* orthologs, respectively (**Extended Data Table 1**). Several members of these OGs are specifically expressed in the endosperm of each angiosperm species and they indeed fall in the same clades (indicated by stars in **Fig. 2d-e**). Remarkably, several genes belonging to these OGs have already been described as PEGs in angiosperm endosperms (indicated by male symbols in **Fig. 2d-e**). Importantly, our study of early-diverging angiosperm transcriptomes also supports an ancestral role for auxin in angiosperm endosperm development in those taxa^15^: in *Amborella trichopoda* and *N. caerulea*, the *YUC10* orthologs *Atrichopoda_scaffold00209*.*9* and *Nycol*.*A00013*.*1* are specifically expressed in their endosperm tissues (**Fig. 2d-e, Extended Data Table 1**).

As alluded to above, the fertilization of the angiosperm megagametophyte is an innovation of the angiosperms, leading to the formation of a biparental endosperm. In gymnosperms, however, the seed nourishing function is provided by the haploid maternal megagametophyte, which develops independently of fertilization. Strikingly, the development of the mature ovule in the gymnosperm *Ginkgo biloba*, which includes the maturation of the megagametophyte, is tightly linked to auxin activity^36^. However, unlike in angiosperms where auxin responses are triggered by fertilization, in *G. biloba* these processes are linked rather to pollen deposition in the ovule^36^. We thus hypothesized that the development of the megagametophyte into a nourishing structure should be linked to auxin biosynthesis in both lineages of spermatophytes, even if its parent-of-origin would be different. If that is true, then orthologues of the angiosperm endosperm auxin biosynthesis genes should be expressed in the gymnosperm megagametophytes. Based on our previous transcriptomic profiles of *Pinus pinaster* megagametophytes, we found that this indeed was the case. Two genes belonging to the above described OGs (PITA_50706, PITA_32408, PITA_5142) are enriched in the *P. pinaster* megagametophyte, and those genes fall in the same clades as the angiosperm endosperm-specific auxin biosynthesis genes (**Fig. 2d-e, Extended Data Table 1**). This allows us to conclude that auxin production is already a feature of gymnosperm megagametophytes, suggesting that a role for auxin in the development of the nutritive tissues of the seed predates the emergence of the angiosperm clade and the origin of the endosperm proper. Because the gymnosperm seed nourishing structure is purely of maternal origin, this means that auxin biosynthesis genes were under maternal gametophytic control in the gymnosperms, and that this control shifted to the father in the angiosperms. These genes thus became PEGs after the angiosperms split from the gymnosperm ancestor.

We thus propose a model where auxin is an evolutionarily conserved driver of seed development in spermatophytes (**Fig. 2f**): in gymnosperms, maternal auxin biosynthesis and activity is triggered by pollination, while in angiosperms these pathways shifted to paternal control, coupling the development of the seed nourishing structure and accessory tissues to fertilization. This innovative switch in control likely ensures a tighter post-fertilization biparental control over resource allocation to the seeds and likely contributed to the evolutionary success of flowering plants.

## Author contributions

AMFR, JBB and DDF designed the experiments. AMFR performed the experiments. LPS and AF performed the metabolomics analyses. MS and AMFR coded the imprinting inference pipeline. AMFR performed the bioinformatic analyses. AMFR and DDF wrote the manuscript with input from all co-authors.

## Data availability

All data generated is available at NCBI under reference PRJNA1157574.

## Statements

The authors declare that they have no conflict of interest.

## Acknowledgments

We thank Ingo Michael at the Botanical Garden of Potsdam and the gardeners at the Botanical Garden of Berlin (BO) for giving us access to the *Nymphaea, Victoria* and *Schisandra* plants. We thank Arun Sampathkumar for help with setting up the LCM system, Barbara Schmitz for help with the auxin treatments and sample collection, and Mehrshad Shobeiri and Maral Charchinezhadamouei for help with the seed measurements.

