## Supplementary Data for "Genomic imprinting in an early-diverging angiosperm reveals ancient mechanisms for seed initiation in flowering plants"

**Fig. S1**

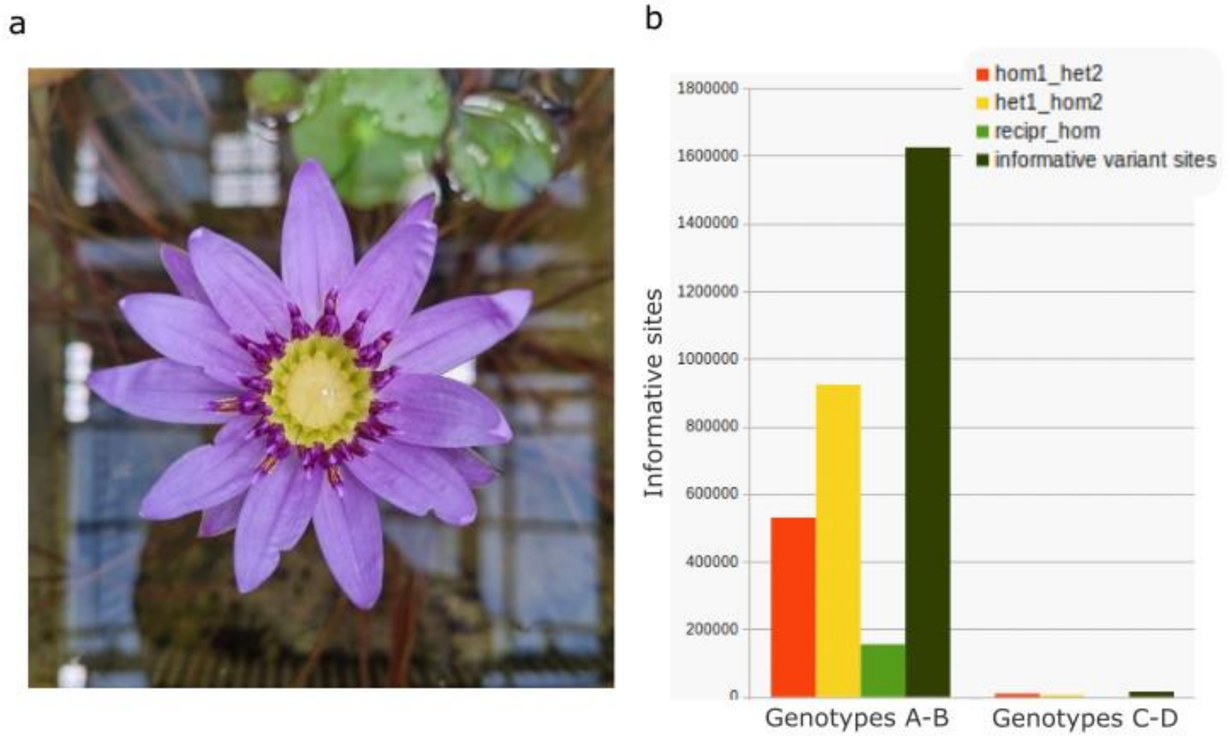


**Extended Data Figure S1**. **a**, Representative photo of a *N. careulea* individual used in this study. **b,** Number of informative sites identified in plants 1 and 2 (here called Genotypes A and B). Genotypes C and D indicate individuals of *Nymphaea colorata* for which there were almost no informative sites. Such individuals are likely clones and are therefore not useful for studies of genomic imprinting. Hom1_het2, sites homozygous in first genotype and heterozygous in the second, het1_hom2, sites heterozygous in first genotype and homozygous in the second, recipr_hom, sites homozygous in both genotypes.

**Fig. S2**

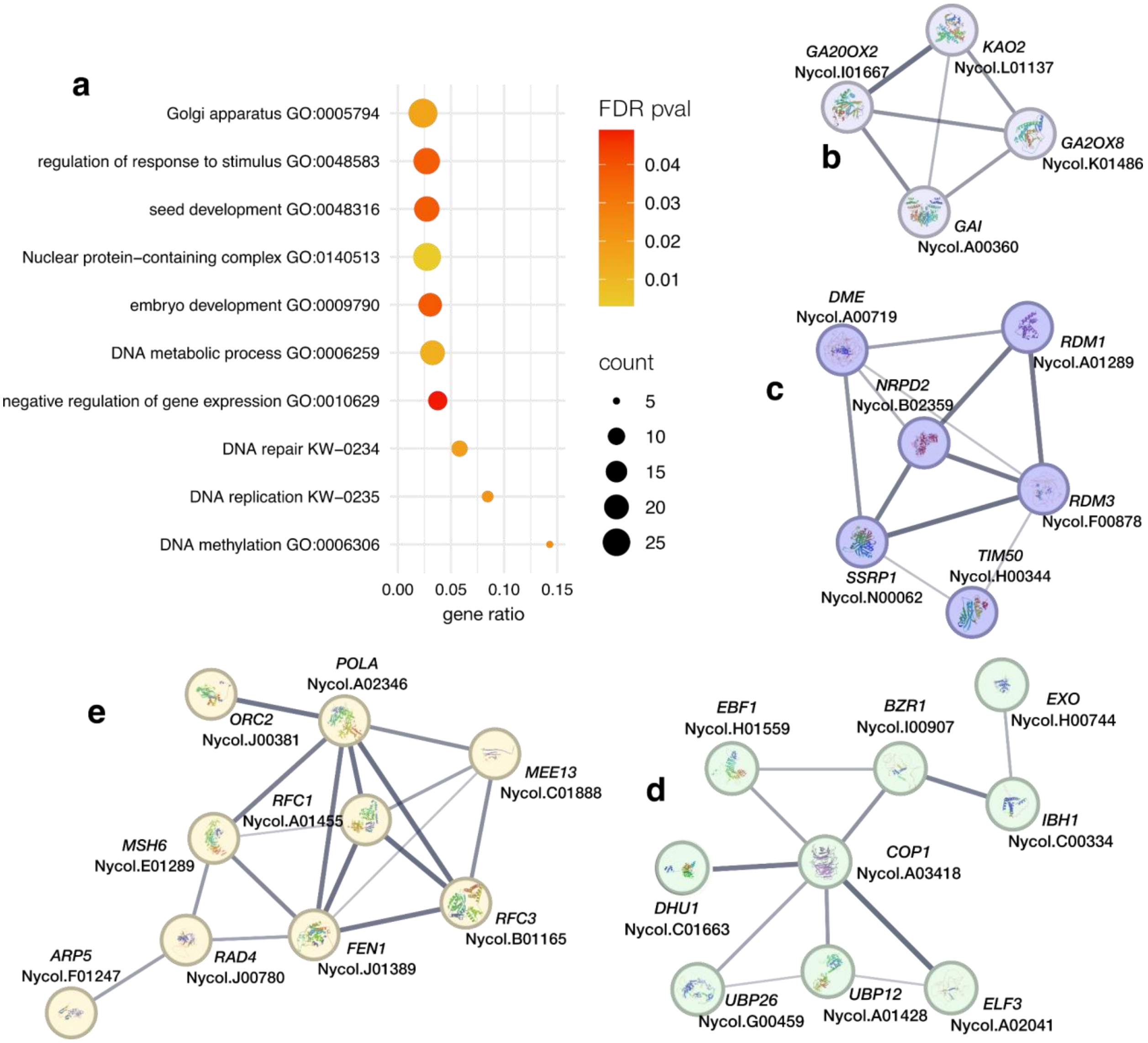


**Extended Data Fig. S2**. **Enriched terms and related putative interaction networks for imprinted genes in *N. caerulea*. a**, Enriched GO categories among genes imprinted in the endosperm of *N. caerulea*. **b**, Gibberellin-related cluster, containing *GIBBERELLIN 20 OXIDASE 2* (*GA20OX2*), *GIBBERELLIN 2-OXIDASE 8* (*GA2OX8*) and *GIBBERELLIC ACID INSENSITIVE* (*GAI*). **c**, Cluster formed by terms “DNA methylation” (GO:0006306; FDR pval 0.0233), “DNA replication” (KW-0235 FDR pval 0.0212) and “DNA repair” (KW-0234 FDR pval 0.0182). These include genes such as: *FLAP ENDONUCLEASE 1* (*FEN1*), *ORIGIN RECOGNITION COMPLEX 2* (*ORC2*), *REPLICATION FACTOR C3* (*RFC3*), *REPLICATION FACTOR C1* (*RFC*) and *DNA POLIMERASE ALPHA SUBUNIT A* (*POLA*), encoding putatively interacting proteins. **d**, **e**, Cluster “negative regulation of gene expression” (GO: 0010629 FDR pval 4.90E-02), including genes such as *RNA-DIRECTED DNA METHYLATION 3* (*RDM3*), R*NA-DIRECTED DNA METHYLATION 1* (*RDM1*), *JUMANJI 25* (*JMJ25*) and *JMJ30*, *ARGONAUTE 1* (*AGO1*) and *AGO10*, and *DNA-DIRECTED RNA POLYMERASE D SUBUNIT*

*2B* (*NRPD2*). These results are in line with observations in Arabidopsis, maize and wild tomatoes, where imprinted genes are enriched for putative chromatin modifying functions or epigenetic regulators and for genes with known functions in seed development (Gehring *et al.*, 2011; Waters *et al.*, 2013; Pignatta *et al.*, 2014; Roth *et al.*, 2018). e Cluster “seed development” (GO: 0048316, FDR pval 3.80E-02), which  included genes such as *OCTOPUS* (*OPS*), *TOPLESS* (*TPL*), *BRASSINAZOLE-RESISTANT 1* (*BZR1*), *DEMETER* (*DME*), *LATE EMBRYOGENESIS ABUNDANT* (*LEA6*), *MATERNAL EFFECT EMBRYO ARREST 13*, (*MEE13*), among other important seed development regulators.
The enriched GO terms and additional details can be found in **Supplementary Table 2**.

**Fig. S3**

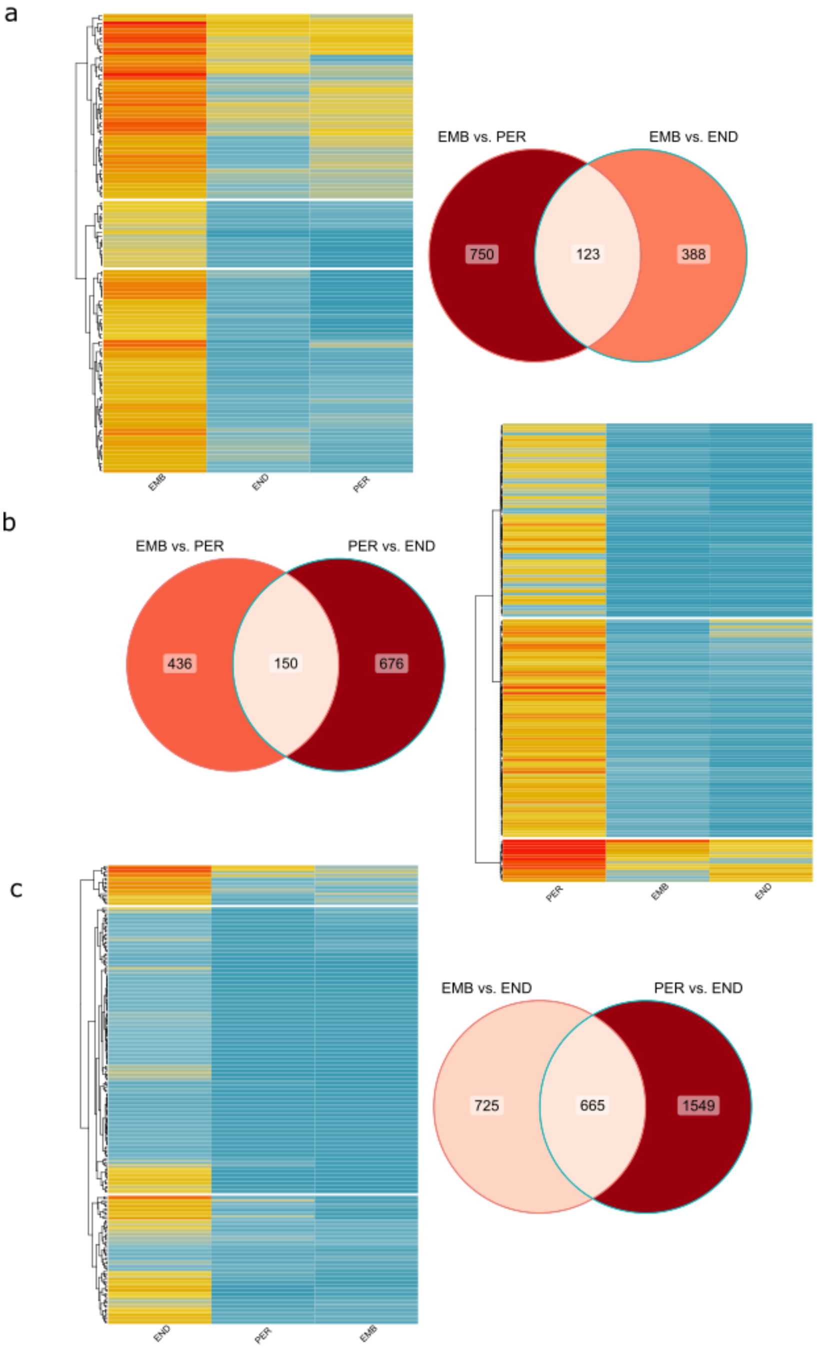


**Extended Data Fig. S3**. **Delimitation of tissue-specific genes based on the intersection of differential expression in pairwise tissue comparisons.** The heatmaps and Venn diagrams show the expression patterns and intersected groups of differentially expressed genes (DEGs) for the following tissues: **a**. 123 embryo, **b**. 150 perisperm, **c**. 665 endosperm-specific genes.

**Fig. S4**

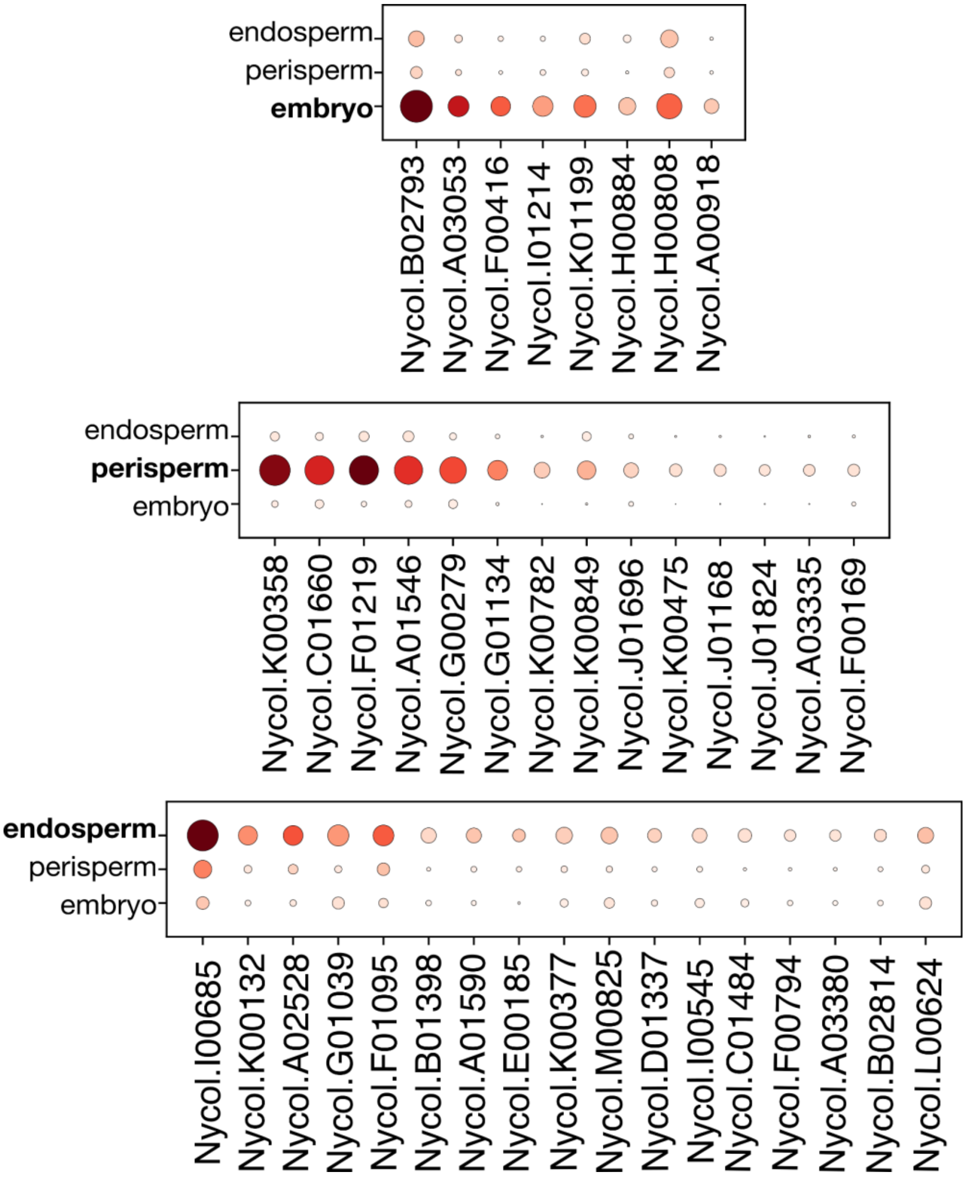


**Extended Data Fig. S4**. **Manual annotation of cell clusters based on LCM expression patterns.**

Dot plots of gene expression of LCM-derived preferentially expressed genes that overlap with the rank genes of the scRNAseq tissue clusters. From top to bottom: embryo, perisperm and endosperm. Darker colors symbolize the mean expression in the cluster and the size of the dot represents the proportion of cells in the cluster in which the gene is expressed. Full data of rank genes per cluster in **Supplementary Table 4**.

**Fig. S5**

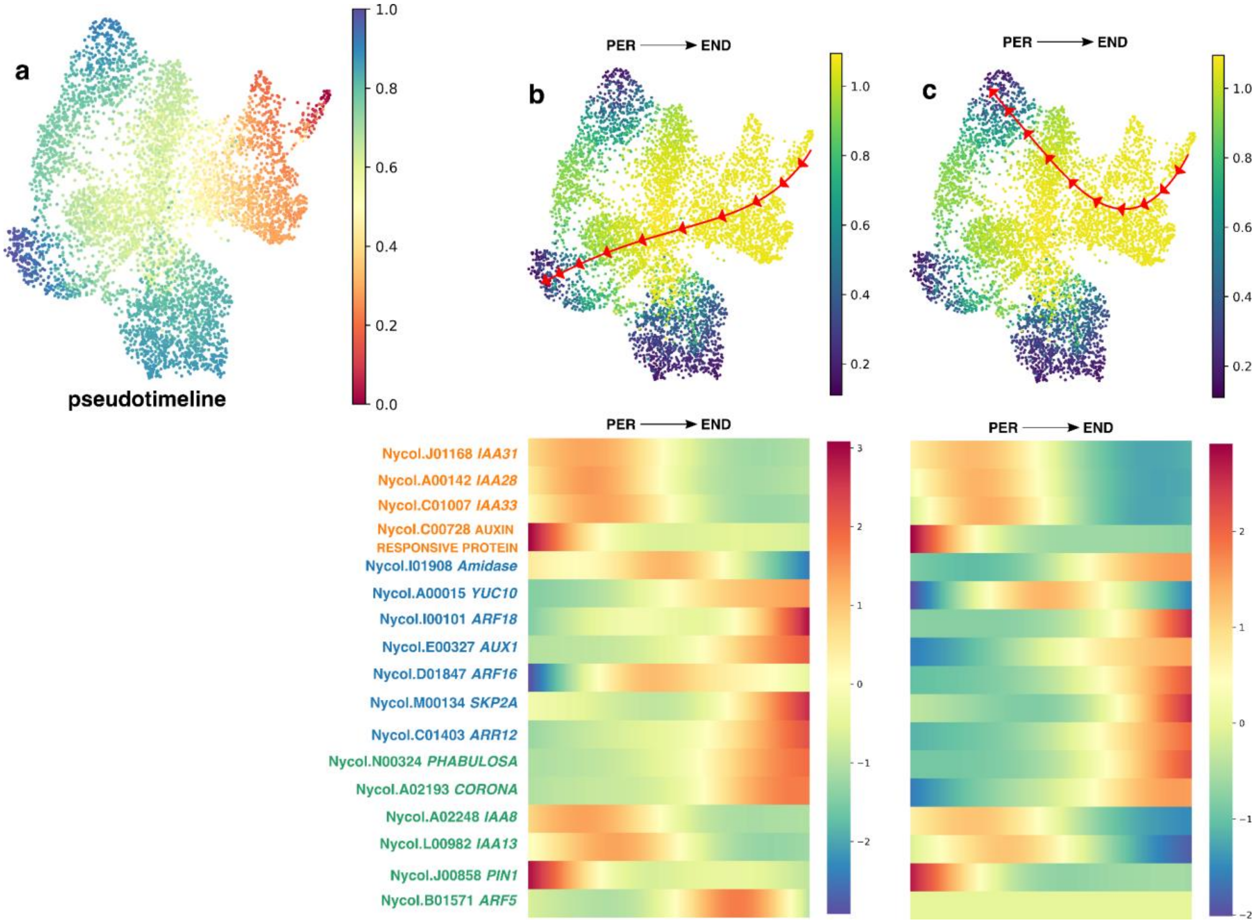


**Extended Data Fig. S5**. **Single-cell trajectory and pseudotime analysis of gene expression in the seed compartments of *N. caerulea*, focusing on auxin-related genes. a**. Pseudotime plot where the red-to-blue color gradient reflects the continuous transition of gene expression states, with red indicating the initial expression profiles and blue marking more advanced changes along the trajectories. **b-c**. Additional trajectories inferred by Palantir, illustrating the transition of gene expression from the perisperm to the endosperm. Dots are colored according to entropy estimates. Below, heatmaps depict the dynamic expression of auxin-related genes across the trajectories.

**Fig. S6**

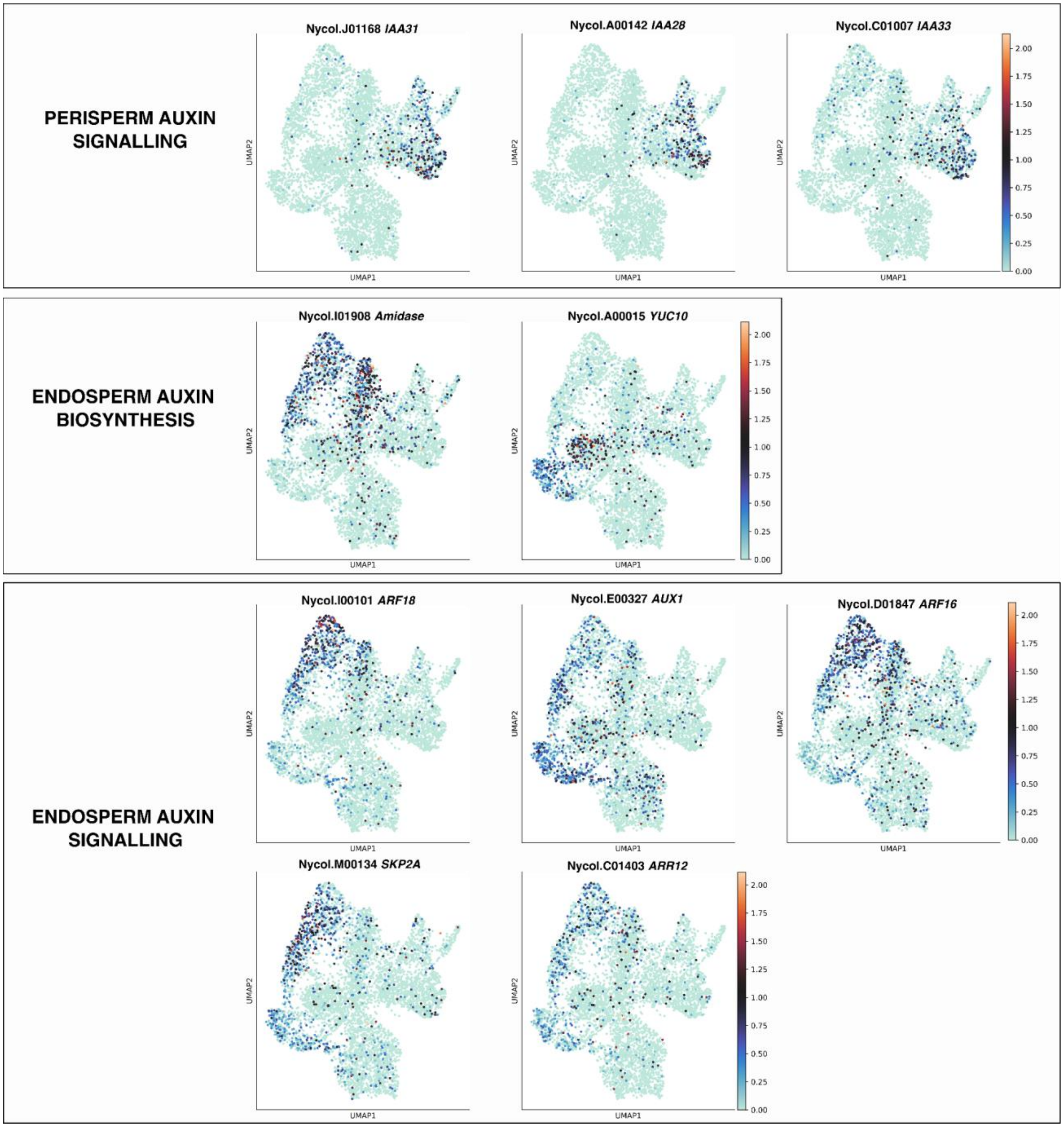
PERISPERM AUXIN SIGNALLING
Nycol.J01168 IAA31
Nycol.A00142 IAA28
Nycol.C01007 IAA33
ENDOSPERM AUXIN BIOSYNTHESIS
Nycol.I01908 Amidase
Nycol.A00015 YUC10
ENDOSPERM AUXIN SIGNALLING
Nycol.I00101 ARF18
Nycol.E00327 AUX1
Nycol.D01847 ARF16
Nycol.M00134 SKP2A
Nycol.C01403 ARR12
UMAP1
UMAP2
0.00
0.25
0.50
0.75
1.00
1.25
1.50
1.75
2.00

### Fig. S6 (cont.)

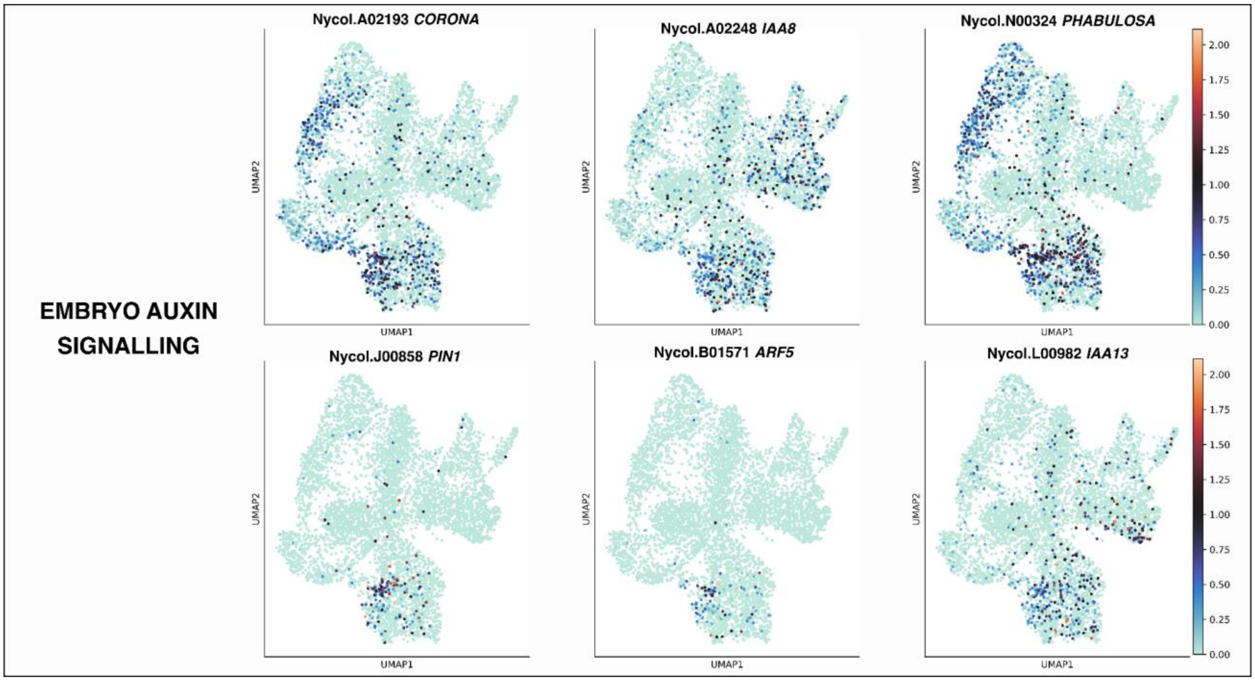


**Extended Data Fig. S6**. **Expression of auxin related genes at the single cell resolution in the *N. caerulea* seed**. UMAP visualization of all nuclei sequenced color coded by levels of expression, from low -cold colors- to high -warm colors–. Sets of genes are grouped according to the tissue they are preferentially expressed and this is reflected on the distribution of the expression across the UMAP.

**Fig. S7**

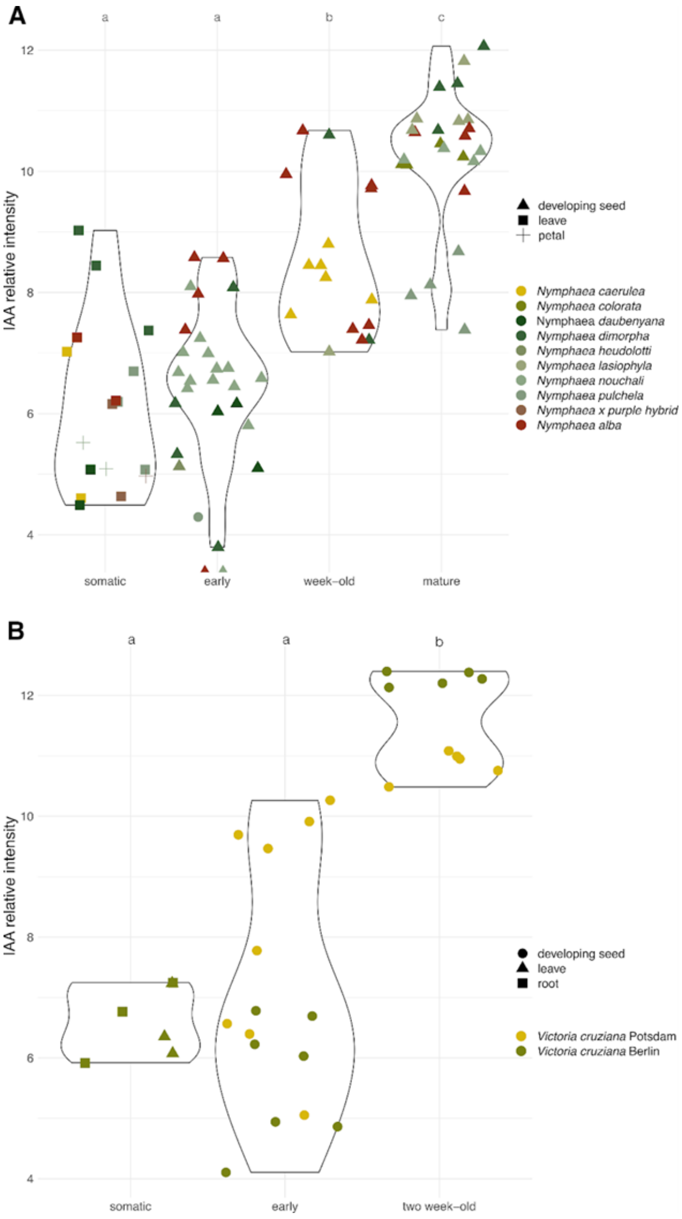


**Extended Data Fig. S7**. Auxin is produced after fertilization in Nymphaeales seeds. **a-b,** Comparative profiling of IAA in somatic tissues and seeds of species of *Nymphaea* (**a**) section Brachyceras, and of *Victoria cruziana* (**b**). IAA levels increase as seed development progresses and its levels in early seeds are similar to those of somatic tissues. Each data point corresponds to a biological replicate (in case of seeds, each datapoint is one seed). Letters indicate Wilcoxon rank sum test, $P < 0.01$.

**Fig. S8**

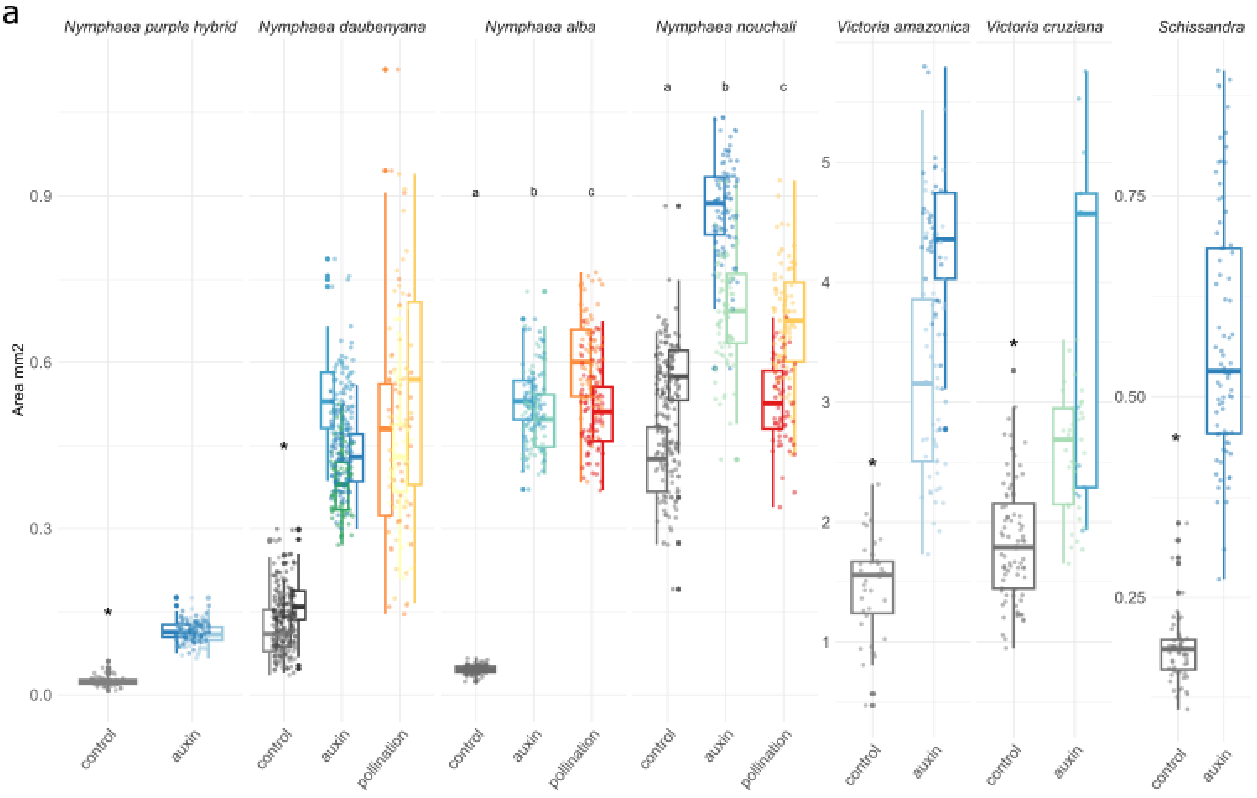


**Extended Data Figure S8**. Auxin experiments in an array of Nymphaeales including species of *Victoria* spp. and in the austrobaileyale *S. chinensis*. Note how exogenous auxin application triggers seed development without fertilization independently of taxa. Mock controls are in black, auxin treatments in cool colors (green-blue) and fertilized seeds in warm colors (yellow-red). Each bar represents one fruit and each dot one seed. Fertilized seeds were not assessed for *N. purple hybrid*, *Victoria* and *Schisandra* due to the lack of compatible fathers. Asterisks and letters indicate Wilcoxon rank sum test, $P < 0.01$.

**Extended Data Table S1. Auxin biosynthesis genes have specialized in the nourishing tissues of spermatophytes**. Differential gene expression summaries of *TAR* and *YUC* ortholog genes that are enriched in the endosperms of angiosperms and in the megagametophytes of *P. pinaster* in comparison to their leaves. Data and analyses as previously published (Florez-Rueda *et al.*, 2024).

| Orthogroup | Locus Name | adj P | logFC | Mean normalized expression leaves | Mean normalized expression nourishing tissue |
|---|---|---|---|---|---|
| OG0001584 - *TAR* orthologs | AT1G23320 | 1,03E-04 | 8,549 | 0,000 | 5,038 |
| | Migut.D01989.1.p | 2,37E-42 | 5,092 | 0,000 | 7,651 |
| | Os05g0169300 | 8,43E-49 | 13,648 | 0,000 | 10,649 |
| | PITA_51242 | 2,01E-98 | 7,750 | 4,678 | 11,156 |
| | Solyc03g112460.3 | 6,69E-211 | 4,222 | 8,357 | 12,572 |
| | Zm00001eb123040 | 1,93E-11 | 4,719 | 2,572 | 7,082 |
| | Zm00001eb281900 | 1,93E-115 | 18,952 | 0,000 | 16,168 |
| OG0001099 - *YUCCA* orthologs | AT1G48910 | 1,31E-26 | 10,574 | 0,000 | 7,102 |
| | Atricophoda_scaffold00209.9 | 4,98E-03 | 10,136 | 0,123 | 4,174 |
| | Nycol.A00013.1.p | 3,17E-08 | 16,936 | 1,770 | 5,847 |
| | Os01g0273800 | 2,02E-25 | 5,638 | 6,620 | 12,294 |
| | Os02g0272200 | 2,95E-14 | 9,582 | 0,000 | 6,655 |
| | Os12g0189500 | 1,10E-32 | 10,977 | 0,000 | 8,371 |
| | PITA_32408 | 7,38E-10 | 9,037 | 0,000 | 4,154 |
| | PITA_50706 | 6,36E-37 | 13,060 | 0,000 | 7,833 |
| | Solyc09g073015.1 | 4,05E-17 | 12,506 | 0,000 | 10,154 |
| | Solyc09g074430.5 | 9,72E-92 | 4,971 | 6,247 | 11,187 |
| | Zm00001eb332870 | 1,80E-08 | 3,413 | 5,429 | 8,504 |
| | Zm00001eb396380 | 2,91E-10 | 3,851 | 1,929 | 5,974 |
| | Zm00001eb409250 | 4,96E-58 | 17,093 | 0,000 | 14,172 |

**Online Methods**

Plant materials

*Nymphaea* plants were screened in the Botanical Gardens of Potsdam and Berlin, Germany, in the summer of 2021 to identify self-incompatible taxa. Several species were screened for diploidy and self-incompatibility. Controlled crosses, emasculations and continuous observations led us to select two polymorphic individuals of the species *N. caerulea* maintained in the living collections of the Botanical Garden of Potsdam. Flowers were bagged before anthesis and seed production was null or low, which is indicative of self-incompatibility. This was in strong contrast to other species of the genus which develop seeds even after emasculations on the day of female receptibility. *N. caerulea* was one of the few *Nymphaea* species that we analyzed where emasculation easily prevented self-fertilization and in which events of self-fertilized seeds were rare in the greenhouse conditions in Potsdam.

Since we aimed to detect imprinting, an important criterion was the levels of polymorphism between accessions, we aimed to have at least two individuals that had polymorphisms between them that allowed for allelic specific expression analyses and imprinting inference. We therefore screen three individuals of *Nymphaea colorata* and two of *Nymphaea caerulea* from the Berlin and Potsdam Botanical Gardens. We collected leaves and got their genomes resequenced, we checked the number of polymorphic sites between them (Figure S1). These genome resequencing datasets were used for imprinting inference, see below. Experiments on *S. chinensis* and *Victoria sp.* were performed using individuals at the Berlin and Potsdam Botanical Gardens.

Hormonal treatments

To test the effect of auxin as an inducer of seed development, we treated a number of Nymphaeales species with 2,4-Dichlorophenoxyacetic acid (2,4-D), a synthetic auxin. The experiments were carried out in the botanical gardens of Berlin and Potsdam. During the spring-summer season, flowers of different Nymphaeales species, namely *N. caerulea*, *N. daubenyana*, *N. nouchali*, *N. alba*, *V. amazonica* and *V. cruziana*, were used for the experiments. We also used a sterile hybrid accession that flowers profusely but never develops seeds after pollination. This accession is called *Nymphaea* x *purple hybrid*.

For the experiments, we identified flowers on their first day of opening when they were female receptive, a state characterized by the accumulation of fluid in the carpels. For negative control experiments we used female receptive plants and carefully removed their anthers, which should be pollen free, and bagged the flowers to prevent pollination in the greenhouse. For pollination experiments, the flowers were pollinated on female receptive days as a positive control. For the auxin application experiments, flowers were emasculated and then 100 μM 2,4-D was applied to the carpel fluid on female receptive days. The volume of 2,4-D solution was adapted to the size of the flower. For example, up to 5 mL to fill the large carpel cup of *Victoria* spp. flowers to a minimum of 100 μL to fill the small carpel cup of *N. alba* flowers.

Further experiments were carried out at the Botanic Garden in Berlin on plants of *S. chinensis* as another representative of an early divergent angiosperm clade, Austrobaileyales. As *S. chinensis* plants are dioecious and only one female individual exists in the collection of the Berlin Botanical Garden, only control and 2,4-D experiments were carried out. This was also the case for the flowers of the *Victoria* spp, as pollinations were not successful in any of our experiments in 2021. Similarly, our pollination experiments with the *Nymphaea* x *purple hybrid* failed to produce viable seeds.

For all experiments in the season of 2021, flowers were collected at 8 DAT (days after anthesis) and immediately fixed by vacuum infiltration in farmer's fixative (9:1, ethanol:acetic acid). The material was kept in the fixative until further processing as described below.

A set of *N. caerulea* plants was then established in the greenhouse of the Max Planck Institute of Molecular Plant Physiology, Potsdam-Golm, in 2022. Further experiments were carried out on these two plants throughout 2022 and 2023 (referred to as plants 1 and 2). We expanded our sample collections to an earlier time point, 3 DAT, and also continued our experiments at 8 DAT. With the aim of identifying a threshold for the effect of auxin, we performed a gradient experiment of increasing auxin concentration from 10 μM to 100 μM 2,4-D. In addition, to test the effects of auxin depletion in the seed, we performed exogenous applications of the auxin inhibitors L-Kynurenine (L-Kyn)(He *et al.*, 2011) and Yucasin(Kakei *et al.*, 2015) to the carpel fluid. We tested two concentrations of each chemical, 30 μM and 100 μM. In brief, plants were manually pollinated on the morning of first flower opening and at least three hours later each inhibitor was applied to the carpel cup. For this set of experiments, whole flowers were collected and frozen at -80 ºC. A piece of the carpel was fixed and processed for histological imaging as described below. For seed size measurements, the seeds were manually dissected from their embedding tissue and carefully separated to allow seed area measurements. Seeds or ovules were placed in Petri dishes with cold PBS 1X, imaged and their area immediately measured using a Keyence digital microscope. For all hormonal treatments mock controls were run in parallel.

<u>Cytological observations</u>

Fruits were manually sectioned to allow embedding in paraffin blocks. The blocks were sectioned at 7 μm on a microtome. Paraffin ribbons were placed on microscope slides, which were further prepared and stained as follows. Slides were dewaxed by immersion in xylol or histoclear twice for 10 minutes. Sections were hydrated through a graded ethanol series to 100% water. For general tissue structure, sections were stained with

1% Safranin for 5 minutes and with Aniline Blue 2% in 3% acetic acid for 1 minute. After a brief water wash, the slides were passed rapidly through a graded ethanol series up to 100% ethanol, taking care to preserve the staining. Finally, the slides were mounted in Roti Mount Aqua. Images were captured on an Olympus motorized epi-fluorescence microscope using 'CellSens' software, and seed area was measured by manually demarcating the most central section of individual seeds using Fiji.

Compartment specific transcriptome generation and LCM

As part of the large set of experiments of 2021, pollinated flowers of *N. caerulea* at one week after pollination were selected for targeted LCM experiments. We aimed to produce transcriptomes of the main compartments of the Nymphaea seed: Endosperm, embryo, and perisperm. We used three different flowers numbered 47, 105 and 115. Flowers were collected from the plants of the Botanical Garden Potsdam a week after pollination. These plants were later established on the MPI-MP in 2022 and used for imprinting inference (see below). LCM was performed as follows. Developing fruits were harvested at 8 DAP after manual pollination. Material was immediately immersed in cold farmers' fixative (Ethanol: Acetic acid solution 9:1) and vacuum infiltrated. The material was kept in the fixative until imbedding which took place in less than a month after collection. Fruits were manually sectioned to allow accommodation in the paraffin blocks. Blocks were sectioned on a microtome at 7 um. LCM protocols were followed as described in (Florez-Rueda *et al.*, 2016a). In brief, capture was performed using a Leica LCM microscope carefully separating the endosperm from the embryo and surrounding perisperm and seed coat tissue. Target tissue was laser captured using a Leica LMD600 microscope at the MPIMP and RNA extraction was performed immediately or within 24 h; in the latter case, the caps were stored at –80 $^{o}$C prior to extraction. RNA extraction was performed using the Applied Biosystems® Arcturus®PicoPure® RNA Isolation Kit (ref. KIT0204, Thermo Fisher Scientific, MA, USA) according to the manufacturer’s instructions. The quality and quantity of total RNA was assessed with Bioanalyzer Pico Chips (Agilent, USA). RNA that showed clear ribosomal peaks was used for further steps. From the extracted RNA, cDNA was amplified using the SMART-Seq® v4 Ultra® Low Input RNA Kit for Sequencing (Clontech Laboratories Inc., USA). cDNA amplification was validated by running Agilent 2100 Bioanalyzer High sensitivity DNA Chip (Agilent, USA). After validating successful library preparation, libraries were sequenced on an Illumina Novaseq 6000 (Illumina Inc., USA) platform at Novogene in the UK.

Crossing experiments for imprinting inference

Reciprocal crosses were performed on the two polymorphic individuals of *N. caerulea* established in the MPIMP during the summer of 2022. Three different flowers per direction of the cross were used for the study and were collected at 7 days after pollination (DAP). For these six samples endosperm tissue was laser captured and its transcriptome was generated as described above. Additionally in 2022, parental genomes were sequenced, DNA was extracted from fresh leaf tissue with the quick DNA miniprep plus kit from (Zymo Research) and DNA was sequenced at the Beijing Genomics Institute.

Bioinformatic analyses of LCM data

Quality of the reads was assessed with Fastqc. Raw reads were filtered by quality, adapters removed and trimmed with Trimmomatic using the parameters: LEADING:3 TRAILING:3 MINLEN:50 HEADCROP:20. We used a Snakemake pipeline to perform further bioinformatic analyses, from mapping up to maternal proportions per gene (https://github.com/mscharmann/ASAP). Mapping of the reads was done against the *N. colorata* v1.2 genome, available in (https://data.jgi.doe.gov/refine-download/phytozome?organism=Ncolorata&expanded=566) using BWA. Variant calling was performed with vcftools (Danecek *et al.*, 2011). Our allelic specific expression pipeline to detect imprinting in the endosperm followed the rationale to not only make use of reciprocal homozygote sites but also of heterozygote sites in either parent as implemented before for the tomato imprintome (Florez-Rueda *et al.*, 2016b). In brief, variant sites between the parental plants were recovered using genome resequencing of both parental individuals. Using a custom Python program, we transformed the frequencies of the discriminant alleles in each SNP into maternal proportions per gene in each direction of the cross. Once a per-gene value was obtained for each direction of the cross, we used thresholds for maternal proportions in reciprocal crosses to call a given gene as potentially imprinted and consider moderately and strongly imprinted genes, as follows: moderate MEGs > 0.75, strong MEGs > 0.9, moderate PEGs < 0.25, and strong PEGs < 0.1 maternal proportion. These thresholds are reciprocally symmetric and consistent with the expectation of a 0.5 maternal proportion of gene expression in the diploid endosperm of *Nymphaea*. Furthermore, genes considered as candidate MEGs and PEGs exhibited significant departures from the expected 0.5, as assessed by chi-square tests with False Discovery Rate corrections. A full list of *N. caerulea* imprinted genes can be found in **Supplementary Table 1**, and the corresponding summary statistics in **Supplementary Table 2**.

Differential gene expression test analyses were performed using a compound factor (tissue) with multiple levels (sample ID) using Deseq2 (Love *et al.*, 2014) in R. For these analyses, the endosperm libraries generated for imprinting inference in 2022 were also included in the analyses. We tested for differential expression between pairs of seed compartments, namely: Endosperm and Embryo, Endosperm and Perisperm, and Embryo and Perisperm. DEGs were defined as those with FDR-corrected p-values < 0.05 and absolute fold changes greater than 2. To delimit a set of genes preferentially expressed in each of the major seed organs, embryo, endosperm and perisperm, we

identified the top set of genes expressed in a given tissue as the intersection of those significantly up-regulated in both comparisons including the tissue. For visualization, Venn diagrams were generated using the package ggVennDiagram (Gao *et al.*, 2021) in R. Full supplementary tables of the total DEGs per comparison and the top sets of DEGs in each tissue are provided in **Supplementary Table 3**.
To detect enriched functions, analyses were performed in STRING (Szklarczyk *et al.*, 2019) and Panther (Mi *et al.*, 2019) for the sets of organ-specific marker genes collected from LCM experiments and for the set of endosperm imprinted genes. All enriched terms reported are significant with an FDR corrected p-value <0.05. To delimit genes with discrete functions, an MCL clustering algorithm with an inflation parameter of 1.5 was implemented in STRING. To describe the expression patterns of marker genes, heat maps were generated using the complexHeatmap package (Gu, 2022) in R.
Lists of imprinted genes in other taxa were checked for overlap with the list of imprinted genes in the *Nymphaea caerulea* endosperm reported here. Lists from *A. thaliana* (Hsieh *et al.*, 2011; Gehring *et al.*, 2011), *A. lyrata* (Klosinska *et al.*, 2016), *Z. mays* (Waters *et al.*, 2013), *O. sativa* (Chen *et al.*, 2018) , *S. bicolor* (Zhang *et al.*, 2016), *L. utitatisimum* (Jiang *et al.*, 2021), *M. lutteus* (Kinser *et al.*, 2021), species of wild tomatoes *Solanum* spp (Roth *et al.*, 2018), and an interspecific cross of the water lilies *N. thermarum* and *N. dimorpha* (Povilus *et al.*, 2024) were collated and checked for overlap.

<u>Nuclei extraction for single cell RNA sequencing</u>

To further investigate the seeds of *Nymphaea caerulea*, we conducted single-cell transcriptomic analysis of seed compartments. Seeds were collected one week (7 days) after manual pollination, by which time they had turned red, indicating successful pollination. We removed the outer red cell layer and dissected the seeds to discard excess starch from the outer perisperm layers. The endosperms, embryos, and surrounding perisperm tissue were then isolated. Up to 150 seeds were manually dissected with at least four washes in fresh, cold PBS 1% to eliminate debris. Nuclei were extracted from the dissected tissue using the CyStain™ UV Precise P kit, following the manufacturer's protocol with minor modifications. All subsequent steps were performed readily, on ice, and reagents were kept cold. The nuclei extraction buffer was supplemented with protease inhibitors, 2% BSA, and RNAse inhibitors (Ambion RNase Inhibitor, SUPERaseIn™ RNase Inhibitor, and RiboLock). The tissue was macerated in the supplemented nuclei extraction buffer for no more than two minutes and immediately filtered through a 30 µm filter. This was followed by centrifugation at 500 xg for 5 minutes at 4°C. After removing the supernatant, 500 µL of staining buffer was added, and the nuclei were gently resuspended before passing through a 10 µm filter. A second centrifugation was performed, and the nuclear solution was reduced to 100 µL. Nuclei were counted using a Neubauer chamber. Library preparation was conducted using 10X Genomics kits according to the manufacturer's instructions, and sequencing was performed by Novogene. Three libraries were successfully generated from the manually dissected *Nymphaea caerulea* seeds.

Bioinformatic analyses of scRNAseq data

Raw sequencing data were processed using Salmon and Alevin-fry (He *et al.*, 2022) to generate gene-level count matrices. Quality control and filtering were conducted with Scampy (Wolf *et al.*, 2018). To address cell-free RNA contamination, we utilized EmptyDrops (Lun *et al.*, 2019) and further corrected for ambient RNA with SoupX (Young & Behjati, 2020). scDblFinder was employed to remove doublets (Germain *et al.*, 2022). Data from three libraries were integrated using Harmony to correct for batch effects (Korsunsky *et al.*, 2019). Clustering was performed with the Leiden algorithm (resolution 0.01), yielding three clusters. Top-ranked marker genes for each cluster were identified (Supplementary Table: Cluster Description). Functional enrichment and network analyses were conducted using STRING (Szklarczyk *et al.*, 2017). Pseudotime and cell trajectory inferences were made using Palantir (Setty *et al.*, 2019) to explore dynamic gene expression changes and lineage relationships.

Metabolomic experiments

Collection of seeds for hormone measurements through mass spectrometry (MS) was performed in the Botanical Gardens of Potsdam and Berlin. For the genus Victoria, two plants of *V. cruziana* were used for the experiments, one in Potsdam and one in Berlin. Plants were cross pollinated but the success of the crossings was very low. Successful pollinations were evident because of increased size of the seeds as they developed. Flowers were collected at an early time point, up to three days after they had opened and when their receptivity was finished. A later time point at two weeks was achieved only twice and only a few seeds were collected for MS measurements. Seeds were carefully dissected and immediately flash frozen and stored for processing at -80 $^{o}$C. Additionally, leaf and root tissue was sampled. Data points correspond to single seeds in each of the plants used.

For the *Nymphaea* seed hormone measurements, a wide array of species of the Nymphaea Section Brachyceras from the Botanical Garden Berlin was used for experiments in 2021. Most accessions available were self-compatible and developed seeds without the need for manual pollination, but manual pollinations were implemented on flowers destined to be used for measurements. Visits were performed weekly and three timepoints of collection were defined: first, an early time point being up to three days after anthesis; second, a week after controlled pollination, characterized by a red color in the seeds in most of the species studied; and third, a later time point at three weeks after pollination, a timepoint in which the seeds are reaching maturity and are colored black or dark brown. Seeds or leaf tissue were dissected and flash frozen and stored for processing at -80 $^{o}$C. Data points refer to independent replicates of pooled seeds per species.

We implemented a liquid:liquid metabolite extraction protocol that allowed for the qualitative analyses of auxin and some of its conjugates (Salem *et al.*, 2020). In brief, extraction was performed using a methyl-tert-butyl-ether (MTBE):methanol (MeOH) solution. The use of acidified water enabled the purification of the phytohormones into the organic MTBE:MeOH phase for further quantification through UHPLC-MS/MS. The quantification of auxin and some of its conjugates was performed with a previously described UHPLC-MS/MS targeted method(Salem *et al.*, 2020) ref. In brief, extraction of the samples was performed using 1mL of a methyl-tert-butyl-ether (MTBE):methanol (MeOH) solution (3:1, v:v) followed by a liquid:liquid purification step with 0.5 mL of water acidified with 0.1% HCl. An aliquot of 500 µL of the upper organic phase containing the phytohoromones was dried and resuspended in 50 µL of MeOH:$H_2O$ (1:1, v:v). The samples were analyzed in a UHPLC-MS/MS system consisting of a Waters® nanoAcquity UPLC® and a Sciex® linear ion trap 4000 QTRAP®. The chromatographic stationary phase consisted of a C18-column (HSS T3 100 mm × 2.1 mm, 1.8 µm diameter particles; Waters®) fitted with a 2.1 × 5 mm guard column (ACQUITY UPLC HSS T3 VanGuard Pre-column, 100 Å, 1.8 µm; Waters®). The mobile phase consisted of a binary solvent system of water containing 0.1% (v/v) formic acid (solvent A) and methanol containing 0.1% (v/v) formic acid (solvent B). The elution gradient were as follows: 62% eluent A for 6.5 min; 45% eluent B from 6.5 to 7.0 min; 10% eluent A from 7.0 to 7.1 min, held at 0% eluent A from 7.1 to 8.1 min and returned to initial conditions by 8.1 min. From 8.1 to 10.0 min the column was re-equilibrated and conditioned to 62% eluent A. The ionization was achieved using electrospray ionization (ESI) through a Sciex® Turbo V® Ion Source operating in negative ionization mode. The mass spectrometer analyzer was operated in multiple reaction monitoring (MRM) method, for the targeted quantification of IAA. The ion transitions (Q1/Q3 Masses: 174.05/130.20 Da) were previously optimized using the authentic standard of IAA. All data processing was performed in the software Analyst 1.6.2 (Sciex®). The relative quantification of IAA across samples was performed based on the total peak area for this compound in each sample.

**Korsunsky I, Millard N, Fan J, Slowikowski K, Zhang F, Wei K, Baglaenko Y,**

**Brenner M, Loh P, Raychaudhuri S**. **2019**. Fast, sensitive and accurate integration of single-cell data with Harmony. *Nature Methods* **16**: 1289–1296.
**Love MI, Huber W, Anders S**. **2014**. Moderated estimation of fold change and dispersion for RNA-seq data with DESeq2. *Genome Biology* **15**: 550.
**Lun ATL, Riesenfeld S, Andrews T, Dao TP, Gomes T, Marioni JC, participants in the 1st Human Cell Atlas Jamboree**. **2019**. EmptyDrops: distinguishing cells from empty droplets in droplet-based single-cell RNA sequencing data. *Genome Biology* **20**: 63.
**Mi H, Muruganujan A, Huang X, Ebert D, Mills C, Guo X, Thomas PD**. **2019**. Protocol Update for large-scale genome and gene function analysis with the PANTHER classification system (v.14.0). *Nature Protocols* **14**: 703–721.
**Pignatta D, Erdmann RM, Scheer E, Picard CL, Bell GW, Gehring M**. **2014**. Natural epigenetic polymorphisms lead to intraspecific variation in Arabidopsis gene imprinting. *eLife* **3**: e03198.
**Povilus RA, Martin CA, Gehring M**. **2024**. Imprinting and DNA methylation in water lily endosperm: implications for seed evolution. : 2024.03.15.585284.
**Roth M, Florez-Rueda AM, Paris M, Städler T**. **2018**. Wild tomato endosperm transcriptomes reveal common roles of genomic imprinting in both nuclear and cellular endosperm. *Plant Journal* **95**: 1084–1101.
**Salem MA, Yoshida T, Perez De Souza L, Alseekh S, Bajdzienko K, Fernie AR, Giavalisco P**. **2020**. An improved extraction method enables the comprehensive analysis of lipids, proteins, metabolites and phytohormones from a single sample of leaf tissue under water-deficit stress. *The Plant Journal* **103**: 1614–1632.
**Setty M, Kiseliovas V, Levine J, Gayoso A, Mazutis L, Pe'er D**. **2019**. Characterization of cell fate probabilities in single-cell data with Palantir. *Nature Biotechnology* **37**: 451–460.
**Szklarczyk D, Gable AL, Lyon D, Junge A, Wyder S, Huerta-Cepas J, Simonovic M, Doncheva NT, Morris JH, Bork P, *et al.*2019**. STRING v11: protein–protein association networks with increased coverage, supporting functional discovery in genome-wide experimental datasets. *Nucleic Acids Research* **47**: D607–D613.
**Szklarczyk D, Morris JH, Cook H, Kuhn M, Wyder S, Simonovic M, Santos A, Doncheva NT, Roth A, Bork P, *et al.*2017**. The STRING database in 2017: Quality-controlled protein-protein association networks, made broadly accessible. *Nucleic Acids Research* **45**: D362–D368.
**Waters AJ, Bilinski P, Eichten SR, Vaughn MW, Ross-Ibarra J, Gehring M, Springer NM**. **2013**. Comprehensive analysis of imprinted genes in maize reveals allelic variation for imprinting and limited conservation with other species. *Proc Natl Acad Sci U S A* **110**: 19639–19644.
**Wolf FA, Angerer P, Theis FJ**. **2018**. SCANPY: large-scale single-cell gene expression data analysis. *Genome Biology* **19**: 15.
**Young MD, Behjati S**. **2020**. SoupX removes ambient RNA contamination from droplet-based single-cell RNA sequencing data. *GigaScience* **9**: giaa151.
**Zhang M, Li N, He W, Zhang H, Yang W, Liu B**. **2016**. Genome-wide screen of genes imprinted in sorghum endosperm, and the roles of allelic differential cytosine methylation. *The Plant Journal* **85**: 424–436.
